# Gut microbiota-derived indole-3 acetic acid mitigates key hallmarks of metabolic dysfunction-associated steatotic liver disease (MASLD) in an NLRP3-dependent manner in *in vitro* and diet-induced mouse models

**DOI:** 10.64898/2026.01.25.701562

**Authors:** Phulwanti Kumari Sharma, Lokesh Kumar, Akash Baghel, Pradipta Jana, Kritida Dhar, Ruby Bansal, Yamini Goswami, Anita Kumari, Rajni Yadav, Shalimar, Shailendra Asthana, Ruchi Tandon

## Abstract

**Background:** Metabolic-dysfunction associated steatotic liver disease(MASLD) is a global epidemic affecting >30 % global population with no approved therapies till date. Recent reports suggest that activation of NLRP3 inflammasome may be involved in the pathogenesis of MASLD. We, therefore, aimed to identify small molecule inhibitor(s) of NLRP3 inflammasome as a potential therapeutic strategy to manage MASLD and validate their efficacy using *in vitro* and *in vivo* models of MASLD.

**Methodology:** We screened an in-house library of natural products using an *in vitro* phenotypic assay and identified a gut microbiota derived metabolite of dietary Tryptophan; indole-3 acetic acid (I3A) for its ability to inhibit the levels of IL-1β and IL-18, which are elevated as a result of activation of NLRP3 inflammasome in differentiated THP1 cells. Subsequently, we carried out several *in vitro* and *in vivo* studies to confirm the mechanism of action of I3A and its ability to mitigate the key hallmarks of MASLD

**Results:** Our *in vitro* data suggest that I3A is an inhibitor of NLRP3 inflammasome. I3A was also found to improve the blood glucose level, plasma lipid profile, hepatic fat, and liver function in high-fat-high-fructose diet induced model of MASLD using C57BL/6 mice.

**Conclusion:** Our results show that I3A, which has been previously reported to be a gut microbiota-derived metabolite of dietary tryptophan, mitigates the key hallmarks of MASLD in an NLRP3 dependent manner. A dedicated structure-activity-relationship (SAR) study around the I3A chemotype may be carried out in future to identify novel NLRP3 inhibitors with desirable pharmacological profile.

## 1. Introduction

Metabolic-dysfunction associated steatotic liver disease(MASLD) is a hepatic manifestation of metabolic disorder [1–2]. It starts with excessive accumulation of lipids in the liver, called as steatosis or NAFL, which further progresses to steatohepatitis (MASH), fibrosis, cirrhosis, and may eventually lead to hepatocellular carcinoma [3–5]. Multiple factors, such as lipid accumulation, oxidative stress, insulin resistance, inflammatory response, and dietary as well as genetic factors, have been proposed as the key factors responsible for the development of MASLD [6–11].

In this regard, the nucleotide binding and oligomerization domain-like receptor family pyrin domain-containing 3 (NLRP3) inflammasome has been reported to contribute to the development of metabolic abnormalities such as insulin resistance, weight gain, and other metabolic complexities, including MASLD [12]. NLRP3, once activated, results in increased expression of caspase-1 and pro-inflammatory cytokines IL-1β and IL-18 [13]. Increased expression levels of NLRP3, caspase-1, IL-1β, and IL-18 have been found in mice fed with a high-fat-high-sugar (HF-HS) diet [14] or a choline-deficient diet [15]. Patients with MASH have also been found to express elevated levels of NLRP3, IL-1β, and IL-18 in the liver biopsy tissues as well as PBMCs compared to controls [16, 17]. NLRP3 knockout mice, on the other hand, were found to be protected from long-term feeding of choline-deficient amino acid-defined (CDAA) diet-induced liver fibrosis [16]. Pharmacological inhibition of NLRP3 has also been found to modulate adiposity, prevent insulin resistance, and reduce liver inflammation and fibrosis in experimental mouse models of MASH [14–15].

We, therefore, aimed to identify small molecule inhibitors of NLRP3, by conducting *in vitro* phenotypic screening of an *in-house* library of small molecules in a cell based assay followed by the detailed pharmacological profiling of hit(s) (Scheme-1). Briefly, from the phenotypic screening, molecules were screened for inhibition of protein levels of IL-1β by ELISA using differentiated THP-1 in the cell culture supernatants followed by inhibition of mRNA levels of IL-1β and IL-18 by RTPCR. In this assay, indole-3 acetic acid (I3A) was found to be a potential inhibitor of NLRP3 inflammasome. Indole-3-acetic acid has been previously reported to be a tryptophan derivative that is produced naturally by the catalytic action of tryptophan mono-oxygenase and indole-3-acetamide hydrolase, found in the gut bacteria [18–19]. Gut microbiota has been previously found to play a crucial role in the regulation of MASLD by synthesizing bacterial metabolites such as short-chain fatty acids, indole derivatives, secondary bile acids, and trimethylamine [20–26]. Interestingly, activation of NLRP3 has also been found to be associated with gut dysbiosis. Mice deficient in NLRP3 have been reported to show altered gut microbiomes such as reduced levels of *Bacteroides* species and enhancement of *Ruminococcus* species [27–28]. There are also some reports suggesting that I3A possesses anti-inflammatory activity in hepatocytes and macrophages and has the capacity to scavenge free radicals [29–30]. Moreover, I3A has also been reported to alleviate high-fat diet (HFD) induced hepatotoxicity via attenuation of hepatic lipogenesis and inflammatory stress in a 12 week study using C57BL/6 mice [31]. A similar study in rats fed with a high-fat diet showed the therapeutic efficacy of I3A in improving MASLD [32]. In spite of these interesting findings, the mechanism of action of I3A has not been reported till date. Subsequent to our finding of I3A being an inhibitor of NLRP3, we conducted an *in vivo* assay to re-confirm the efficacy of I3A in mitigating the key hallmarks of MASLD in a high-fat-high-fructose (HF-HF) diet induced model of MASLD using C57BL/6 mice. This is the first report suggesting that I3A mitigates the key hallmarks of metabolic-dysfunction associated steatotic liver disease in mice in an NLRP3 dependent manner.

**Scheme-1:**
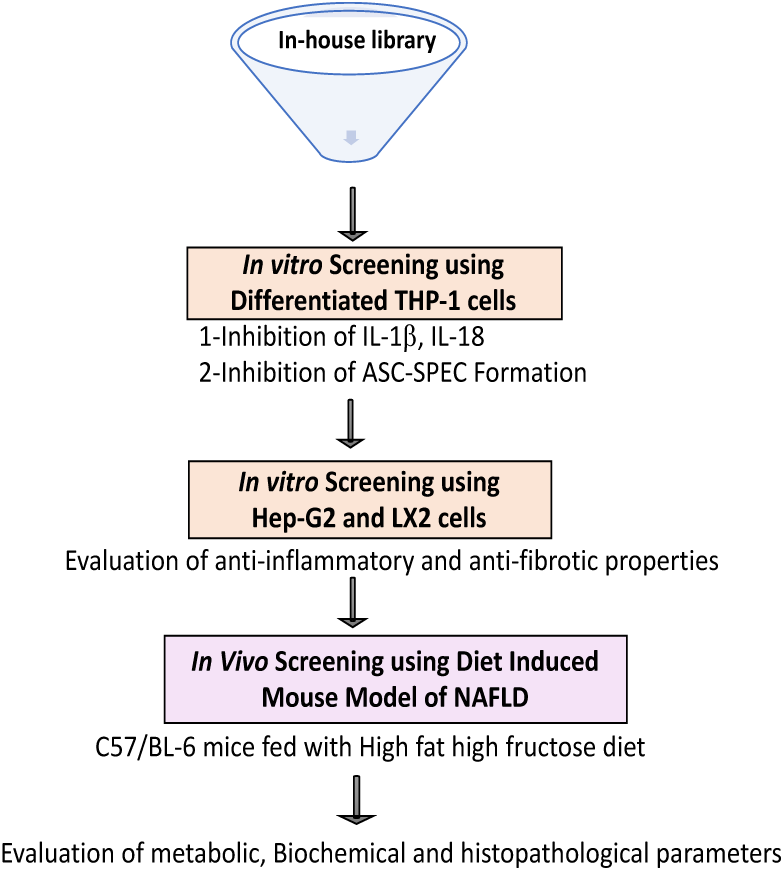
Schematic representation of flow of *in vitro* and *in vivo* experiments.

## 2. Materials and Methods

Following schematic plan was adopted to screen the compounds in the *in vitro* and *in vivo* model of MASLD:

### 2.1 Cell culture

THP-1 and Hep-G2 cells were procured from (American Type Culture Collection) (ATCC) and maintained in RPMI-1640 (Gibco, USA), and DMEM(Gibco, USA) respectively and supplemented with 10% fetal bovine serum (FBS) and 5% penicillin-streptomycin. The cells were incubated at 37 °C in a CO_2_ incubator. No cell line was used above 20 passage number for any of the experiments.

### 2.2 Inflammasome activation analysis

THP-1 cells (passage-5) were seeded at a density of 4 X 10^3^ cells/well in a 96-well plate and differentiated for 48 h with phorbol-12-myristate-13-acetate (50 nM). Cells were then washed thrice with PBS and replenished with RPMI medium containing 10% FBS and 5% penicillin-streptomycin. Cells were then treated with various concentrations of indole-3-acetic acid for 30 minutes and primed with LPS (100ng/ml) for 2 hours, followed by a second stimulation with ATP (2.5 mM) for 2 hours to activate the NLRP3 inflammasome. Cell culture supernatants were used for ELISA and cell lysates were used for RTPCR. For gene expression analysis, cells were lysed in Tri reagent (Sigma), followed by isolation of RNA and preparation of cDNA as per manufacturer’s instructions. Levels of IL-1β and IL-18 were analyzed by RTPCR using gene specific primers for SYBR chemistry on the QuantStudio 6 Flex Real-Time PCR System (Applied Biosystems).

### 2.3 Cell viability assay

To determine the cytotoxicity of I3A, 3-(4,5-dimethylthiazol-2-yl)-2,5-diphenyltetrazolium bromide (MTT) assay was performed. Cells treated as above in the inflammasome activation assay and treated with MTT (100 µl of 0.5 mg/ml MTT stock in PBS) in an independent experiment, and the plate was incubated for 4 h at 37 °C in a CO_2_ incubator. The MTT mixture was aspirated and 100 µl of DMSO was added to each well to dissolve the formazan particles. The plate was shaken for 30 min on a plate shaker at 37 °C to ensure complete dissolution of formazan crystals and read at 570 nm to measure the probable cytotoxicity of test substances if any.

### 2.4 ASC-SPEC formation assay by confocal microscopy

THP-1 cells were plated at a density of 2×10^3^ in an 8-well chamber slide (Genetix Biolabs Asia Pvt. Ltd.) and differentiated with PMA (50 nM) for 48 h. Cells were washed thrice with PBS, replenished with RPMI medium containing 10% FBS and 5% penicillin-streptomycin and treated with the test substances, followed by treatment with compound and inflammasome activation, as stated above. Cells were then fixed with 4% paraformaldehyde for 30 min, blocked with 1% bovine serum albumin for 30 min, and incubated with an anti-human ASC antibody (Santa Cruz Biotechnology Inc., Santa Cruz, CA, USA) at 4° C overnight. Cells were further incubated with an anti-rabbit Alexa Flour 488 antibody (Invitrogen, Carlsbad, CA, USA) for 2 h at room temperature. Cells were co-stained with 4’,6-diamidino-2-phenylindole (DAPI, 2 µg/mL; Invitrogen, Carlsbad, CA, USA) for nuclear staining. Slides were mounted in fluorescent mounting medium (Vector Laboratories, Burlingame, CA, USA), and samples were examined with an FV3000 laser scanning confocal microscope (Olympus, Tokyo, Japan).

### 2.5 3D-Spheroid assay using a co-culture of HepG2 and LX2 cells

A three-dimensional spheroid model was established by co-culturing the hepatocellular carcinoma cell line, HepG2, and the hepatic stellate cell line, LX2, as previously published [33–35]. HepG2 cells (passage-7), were maintained in DMEM (Thermo Fisher Scientific Inc., USA) containing 10% FBS and 1% penicillin-streptomycin, whereas LX2 cells (passage-4) were maintained in DMEM containing 2% FBS and 1% penicillin-streptomycin. To set up the 3D spheroids, a co-culture of HepG2 (2.5×10^3^ cells/well) and LX2 (0.5×10^3^cells/well) cells in DMEM containing 10% FBS and 1% penicillin-streptomycin (Thermo Fisher Scientific Inc., USA) was seeded into the ultra-low attachment, 96-well plate (Corning, USA) and incubated at 37 °C for 48 h to allow the formation of compact, three-dimensional, spheroids in each well. The media was replaced with fresh DMEM, and spheroids were treated with test substances as indicated, followed by treatment with palmitic acid (500 µM) and the plate was incubated at 37 °C in the CO_2_ incubator for an additional 48 h. The plate was centrifuged at 1000 × g (Eppendorf centrifuge 5810R), spheroids were lysed in the lysis buffer, and cDNA was prepared following the manufacturer’s protocol (Thermo Fisher Scientific Inc., USA). RTPCR experiment was done to evaluate the mRNA expression levels of fibrosis marker genes (α-SMA, COL1-A1, and TGF-β), using the TaqMan™ Gene Expression Assay (FAM) (part number 4453320; Applied Biosystems), and 18S was used as an internal control for normalizing the gene expression across the wells using TaqMan™ Gene Expression Assay, VIC (Part number 4448489; Applied Biosystems) on the QuantStudio 6 Flex Real-Time PCR System (Applied Biosystems).

### 2.6 Immunofluorescence studies of 3D-spheroids by confocal microscopy

Spheroids were washed twice with PBS and fixed by incubating them with 4% paraformaldehyde (PFA) at room temperature for 30 minutes. Spheroids were then washed three times with PBS, blocked with 5% horse serum in PBS containing 0.5% triton X-100 for 30 minutes at room temperature, and washed three times with PBS. Organoids were then incubated with primary human HNF4 (Cell Signaling Technology Inc., USA) and primary human COL1A1-Alexa fluor 647 antibody (Cell Signaling Technology Inc., USA) at 1:3000 and 1:100 dilutions, respectively, in the blocking buffer for 90 minutes at room temperature followed by washing three times with PBS, giving 5 minutes for each wash. Spheroids were then incubated with anti-rabbit IgG-Alexa fluor 488 secondary antibody (Invitrogen, USA) at a 4 µg/ml concentration with a 1:1000 dilution in the blocking buffer for 60 minutes at room temperature, followed by washing three times with PBS. Samples were then incubated with DAPI (5 µg/ml in the blocking buffer) for 10 minutes at room temperature for nuclear staining and washed again three times with PBS. A hundred microliters of fresh PBS was then added to the samples, and spheroids were analyzed at 20X magnification by a confocal microscope (Olympus FV 3000).

### 2.7 Computational studies to get the most likely binding pose of I3A

#### 2.7.1 Protein and ligands preparation

The crystal structure of the human NLRP3 NACHT domain (PDB ID: 7ALV) was retrieved from RCSB (Research Collaboratory for Structural Bioinformatics) and was used for the docking studies [36–39]. The protein structures were prepared by using the protein preparation wizard (PPW) module of Maestro (Schrodinger LLC, New York, NY, USA). The systems were pre-processed by adding the hydrogen atom, missing side chains, and loops using the Prime module of the Schrodinger suite [40–41]. Then the structures were prepared with protonation states of residues at neutralizing pH-7 (neutral biological pH) predicted by the PROPKA module in H-bond assignment panel PPW [42–44]. Finally, the hydrogen bond optimization and restrained minimization were done by using the OPLS3 (Optimized Potentials for Liquid Simulations) force field [45–47]. Both molecules were prepared by using the Ligprep module of the Schrodinger suite (Schrodinger, LLC, New York, NY, 2021).

#### 2.7.2 Molecular docking

Docking studies were carried out using the *Glide* module of the Schrodinger suite. The co-crystal ligand “RM5” of NLRP3 is considered a “control” in the docking studies. The RM5 is a close analog of the known control MCC-950, which is another reason for its selection as a control. The grids were generated using the centroid of control molecules (co-crystal) by using the Receptor Grid Generation panel in *Glide* [45, 48]. Based on benchmarking with control, the dockings of indole-3-acetic acid and tryptamine were done. The final docked poses were processed using the Prime MM-GBSA panel in Schrodinger Suite (Schrödinger Suite, LLC, New York, The Y, 2020).

#### 2.7.3 Binding energy analysis

The average binding energy was calculated for a complex by using the PRIME MM-GBSA (molecular mechanics/generalized born surface area) module of the Schrodinger suite. The binding energy was calculated using the following equation:

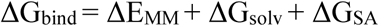

where the difference in the minimized energies between ligand and protein complexes is denoted by ΔEMM. ΔGsolv is the difference in the GBSA solvation energy of the complexes and the sum of solvation energies for the protein and ligand, whereas the differences in surface area energy of the complex and the sum of that in the protein and ligand [39].

### 2.8 Animals

8-12-week-old male C57BL/6 mice (20-25 g) were procured from the Small Animal Facility of the Translational Health Science and Technology Institute (THSTI), Faridabad, and housed in individually ventilated cages with controlled air flow as per the institute’s experimental animal guidelines. The animal room has a controlled 12 h light and 12 h dark cycle with a temperature of 25 ± 2 °C and a relative humidity of 60 ± 10%. The animals were acclimatized for one week in the above conditions prior to initiating the experiments. All the protocols for animal experiments were approved by the Institutional animal ethics committee (IAEC) of THSTI, Faridabad, India (Protocol approval no. IAEC/THSTI/168).

### 2.9 *In vivo* efficacy studies in C57BL/6 mice fed with an HF-HF diet

C57BL/6 mice were divided into five different groups with respect to their initial body weight. The first group (control) was kept on a chow diet, and the remaining four groups were fed with high-fat, high-fructose (HF-HF) diet (Research Diet, USA). Detailed composition of the animal diet is provided in the Table-1. The number of mice in each group was 6 (n=6), and 4 weeks after HF-HF feeding, animals were re-grouped on the basis of weight, fasting blood glucose (FBG), and tri-glyceraldehyde (TG) to initiate treatment with (HF-HF+MCC-950, 20 mg/kg), (HF-HF+I3A, 50 mg/kg b.w), Saroglitazar (Zydus Cadila Healthcare Ltd.) (HF-HF+S, 3 mg/kg b.w.), and HF-HF alone in the control group daily for the next 18 weeks by oral route using 0.5% carboxymethylcellulose (CMC) as a vehicle. Doses of MCC-950, I3A and Saroglitazar in the *in vivo* studies were selected based on the literature evidences [14, 31, 34]. After 18 weeks of treatment, an intraperitoneal glucose tolerance test (IPGTT) was also performed along with their body weight, tail/body length, fasting blood glucose, and lipids. Mice were euthanized, and blood and liver tissues were collected and weighed after washing with chilled phosphate buffer saline (PBS). The plasma and tissue samples were stored at − 80 °C freezer for further analysis.

**Scheme-2:**
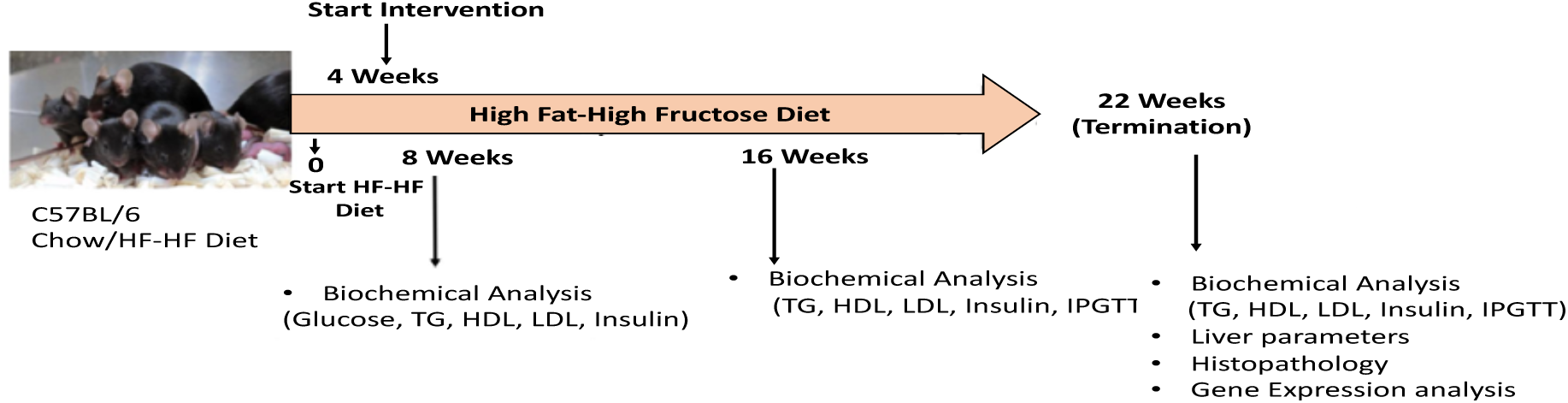
Schematic representation of development of mouse model of non-alcoholic fatty liver diseases in a 22 week study using C57BL/6 mice fed with and high fat-high fructose (HF-HF) diet

#### 2.9.1 Biochemical analysis

Plasma triglycerides (TG, Randox TRIGS kit; Cat. No. TR2774), cholesterol (CHL, Randox CHOL kit; Cat No. CH2773), low-density lipoprotein (LDL, Randox LDL kit; Cat No. CH2776), and high-density lipoprotein (HDL, Randox HDL kit; Cat No.CH2775) were measured in plasma samples using the manufacturer’s protocols.

#### 2.9.2 Intraperitoneal glucose tolerance test (IPGTT)

Animals were fasted for six hours for the IPGTT assessment before and after the treatment period. Glucose was administered (2 g/kg, i.p.) to the fasted mice, and blood glucose concentrations were measured at 0, 15, 30, 60, 90, and 120 min after glucose administration, using a glucometer (Accu-Chek, India).

#### 2.9.3 Histopathological analysis

Small pieces of liver tissue after washing with PBS were fixed in 10% formalin, sectioned, and stained with hematoxylin and eosin (H&E), and Masson’s trichrome (MT) stains. Histological evaluation and MASH-CRN grading and staging were performed at X200 magnification at the All India Institute of Medical Sciences (AIIMS), New Delhi.

#### 2.9.10 Gene expression studies

RNA was isolated from the cells or the mouse liver tissues using the TRI reagent (Sigma Aldrich) following the manufacturer’s protocol. The quality and concentration of the isolated RNA were determined using a NanoDrop spectrophotometer (Thermo Scientific), and 2 µg of RNA was used to synthesize cDNA (Applied Biosystems) as per the manufacturer’s protocol. Real-time Polymerase Chain Reaction (RT-PCR) was carried out on Real-time PCR QS6 (Applied Biosystems) using SYBR Green master mix and the gene-specific primers (Table-2). Gene expression data was normalized against the housekeeping gene, GAPDH.

## Statistical analysis

Each experiment was performed three or more times independently. Statistical analysis was performed on GraphPad PRISM 8.3 software (GraphPad, La Jolla, CA, USA) using one-way ANOVA or two-way ANOVA followed by Bonferroni’s test. The p-values of <0.05 were considered statistically significant. The results were calculated as mean ± SD, and the data are shown as mean + SEM.

## 3 Results

### 3.1 *In vitro* screening of an in-house library of small molecules

The *in house* library of small molecule from natural products was screened in the THP1 cells by looking at the levels of IL-1β as a phenotypic marker of inflammasome activation. PMA differentiated THP-1 cells, when primed with LPS followed by stimulation with ATP, led to increased mRNA levels of IL-1β and IL-18 by RTPCR while pre-treatment of cells with I3A or MCC-950, showed a decrease in levels of IL-1β (Figure 1A). Cell viability was done to ascertain the cellular toxicity of I3A, if any, by MTT method (Figure 1B). We also looked at the mRNA expression levels of IL-1β and IL-18 in this assay wherein pre-treatment of cells with I3A or MCC-950, showed a decrease in the mRNA levels of IL-1β and IL-18 (Figure 1C and 1D).

**Figure 1:**
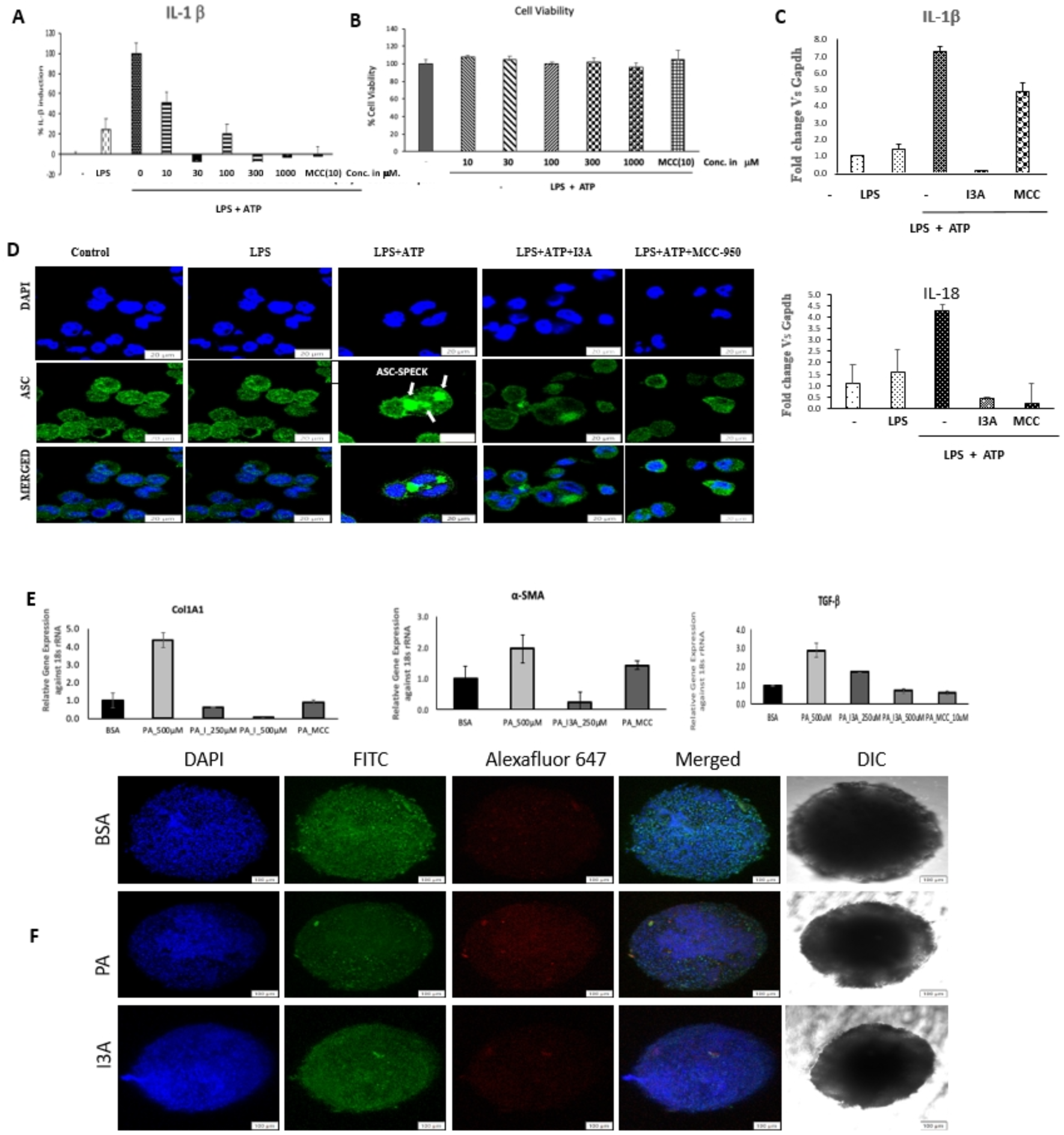
*In vitro* data of I3A-. *In vitro* cell based data in PMA differentiated THP1 cells primed with LPS followed by stimulation with ATP. (A) Inhibition of protein levels of IL-1β by ELISA (B) Cell viability assay by MTT method. (C) Inhibition of mRNA levels of IL-1β and (D) IL18 by I3A(100 μM) and MCC-950(10 μM). (E) Inhibition of ASC-SPEC formation by I3A and MCC-950 compound. *In vitro* 3D spheroid assay using Hep-G2 and LX2 cells to test the anti-fibrotic potential of compounds. (G) Confocal imaging showing the inhibition of COL1A1 staining by I3A; (F) RTPCR assay showing inhibition of palmitic acid induced mRNA expression of and COL1A1, α-SMA and TGF-β by I3A and MCC-950 compounds in 3D spheroids.

### 3.2 I3A inhibits ASC-SPEC formation in THP1 cells

Priming of differentiated THP1 cells with LPS followed by stimulation with ATP led to the formation of a single micrometre size ASC-SPEC in each cell as analyzed by immunofluorescence analysis using confocal microscopy. Pre-treatment of THP-1 cells with I3A or MCC-950, on the other hand, inhibited the LPS+ATP induced SPEC formation in these cells by microscopic analysis (Figure 1E).

### 3.3 I3A inhibits the expression of palmitic acid-induced fibrosis markers in the 3D spheroid assay

The effects of I3A and MCC-950 were analysed for inhibition of palmitic acid (PA) induced fibrosis in the 3D spheroid model using a co-culture of Hep-G2 and LX2 cells [33–34]. Treatment of 3D spheroids with PA (500 µM) led to a significant increase in the mRNA levels of fibrosis markers such as α-SMA, COL1A1, and TGF-β as shown previously [. Both I3A and MCC-950 reduced the expression of PA-induced α-SMA, COL1A1, and TGF-β (Figure 1F). Expression analysis of the hepatocyte marker HNF4 and the fibrotic marker COL1A1 and DAPI for nuclear staining was also performed in the spheroids by immunofluorescence studies using confocal microscopy. Similar to gene expression data, confocal images also showed a reduction in the COL1A1 staining in the I3A-treated spheroids compared to untreated spheroids without compromising the HNF4 or DAPI staining, suggesting an anti-fibrotic activity of I3A (Figure 1G).

### 3.4 In-silico data

The co-crystal structure of the NLRP3-NACHT protein bound with inhibitor RM5 and ADP comprises the four typical subdomains, a nucleotide-binding domain (NBD), helical domain 1 (HD1), winged-helix domain (WHD), and helical domain 2 (HD2) [48] (Figure 2). From the analysis of the co-crystal, the inhibitor (RM5) pocket constituted by residues A227, A228, G229, I230, R351, P352, V353, M408, F410, I411, L413, V414, T439, T524, I574, F575, R578, Q624, S626, L628, E629, Y632, T659, and M661 was identified. These residues are marked by taking a distance of 3.5 Å from the centre of the ligand RM5. From the docking results, we identified that the residues A227, A228; R351; and R578 are key residues, as the ligands established crucial interactions with them in the form of hydrogen bonds. The additional residue is R578 as it supports pi-cationic interactions with ligands.

**Figure 2:**
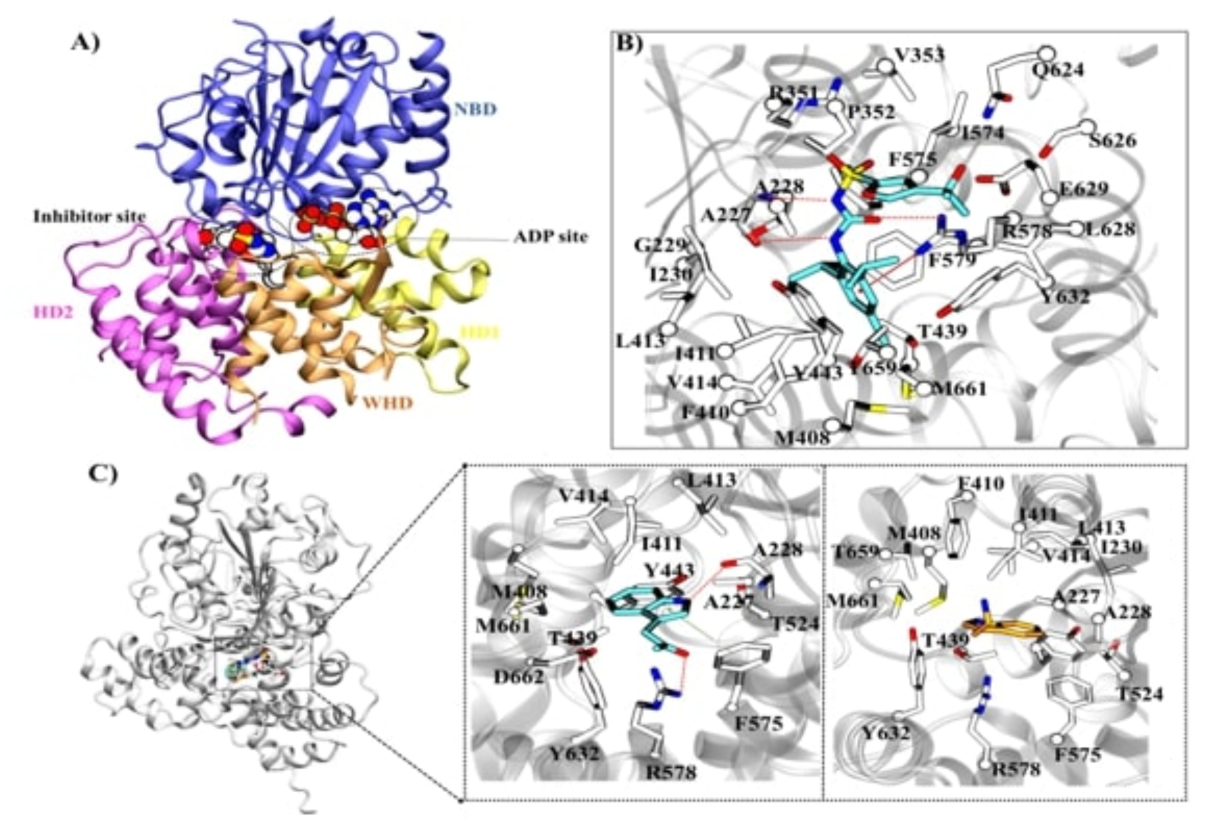
Characterization of NLRP3 and I3A binding site through molecular docking: **A)** Overall crystal structure of human NLRP3 NACHT domain with a subdomain, nucleotide-binding domain NBD (blue), helical domains HD1 (yellow) and HD2 (magenta), and the winged-helical domain WHD (orange). The inhibitor and ADP are shown in vdW. **B**) The interaction diagram of the inhibitor binding site in NLRP3. The red dotted lines and solid line indicate the hydrogen bonding and pi-cation interactions with proteins, respectively. **C)** The superimposition of the ligand with the docked pose of indole 3-acetic acid and tryptamine. The interaction diagram for Indole-3 acetic acid is highlighted. The hydrogen bond and pi-pi interactions are shown in red and green dotted lines.

To optimize the binding pose of the ligand, focused docking was carried out, which is well supported by binding energy calculations on the inhibitor site of NLRP3 with the indole 3-acetic acid (I3A) and tryptamine molecules. The docking and MM-GBSA scores for co-crystal ligand are −8.99 kcal/mol and −71.19 kcal/mol, respectively, while for indole-3 acetic acid and tryptamine they are −5.66, -4.32 kcal/mol, and −25.52, -17.62 kcal/mol, respectively (Figure 2). The superimposition of the docked complex with the crystal structures also revealed that the molecules exhibit a similar crystal binding pose in the pocket. The compound indole-3 acetic acid has shown interactions with the residues A227, A228, I411, L413, V414, Y443, T524, F575, R578, M408, M661, and D668 (Figure 2). The I3A forms hydrogen bonds with the residues A228 and R578 and a pi-pi interaction with residue F575 (Figure 2). It is interesting to underscore that the I3A forms a similar key interaction as found in co-crystal with ligand RM5. However, in the docked complex of tryptamine, it only has hydrophobic and van der Waal interactions with the residues A227, A228, I230, M408, F410, I411, L413, V414, T439, T524, F574, R578, Y632, and M661 (Figure 2)

### 3.5 I3A improves the key metabolic parameters in the HF-HF diet induced model of MASLD

A significant and consistent gain in the overall weight of the animals was seen after 8 weeks of HF-HF diet feeding. Treatment of mice with Saroglitazar (HF-HF+S), MCC-950 (HF-HF+MCC-950), and Indole-3 acetic acid (HF-HF+I3A) for 18 weeks showed a reduction in body weight. A statistically significant reduction in body weight was observed in the HF-HF+S and HF-HF+I3A groups (Figure 3A). Similarly, a significant increase in the body mass index (BMI) was observed in the HF-HF-fed mice in comparison to the control chow-fed mice. The BMI of the Saroglitazar and I3A-treated mice was significantly reduced as compared to the HF-HF group (Figure 3B). In addition to these, mice treated with HF-HF+I3A, and HF-HF+MCC-950 also showed a slight reduction in the liver weight compared to mice in the HF-HF group (Figure 3C).

**Figure 3:**
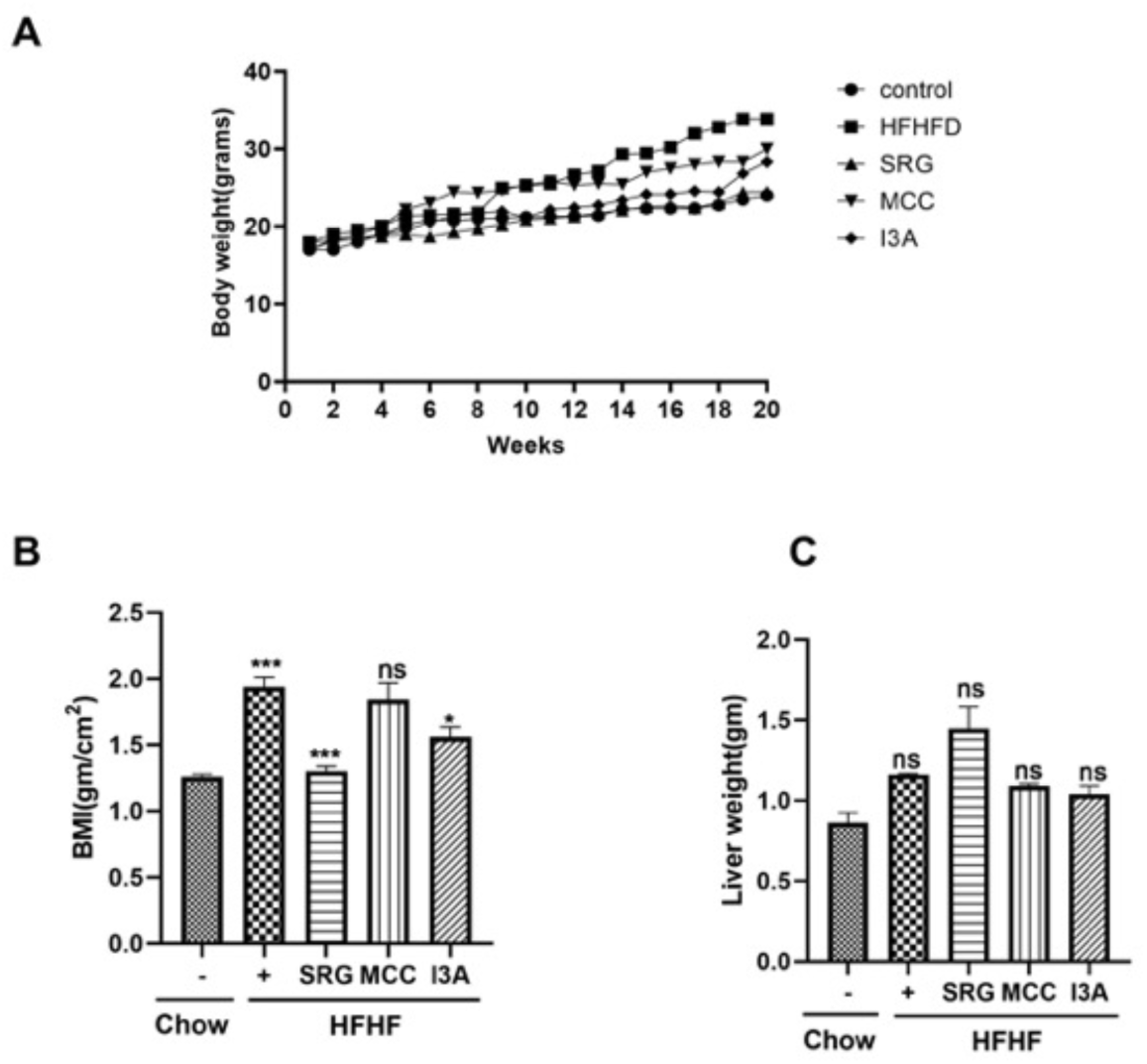
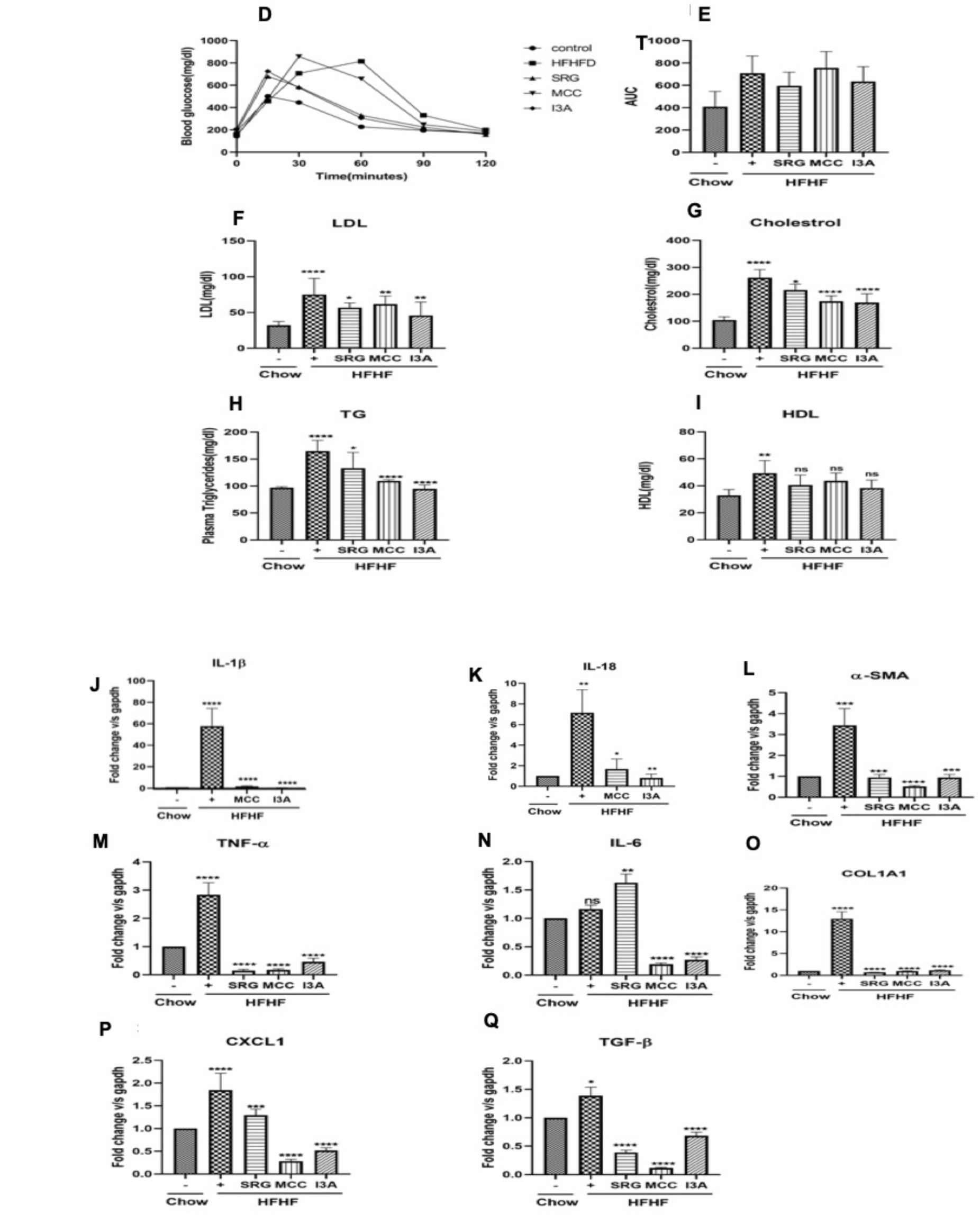
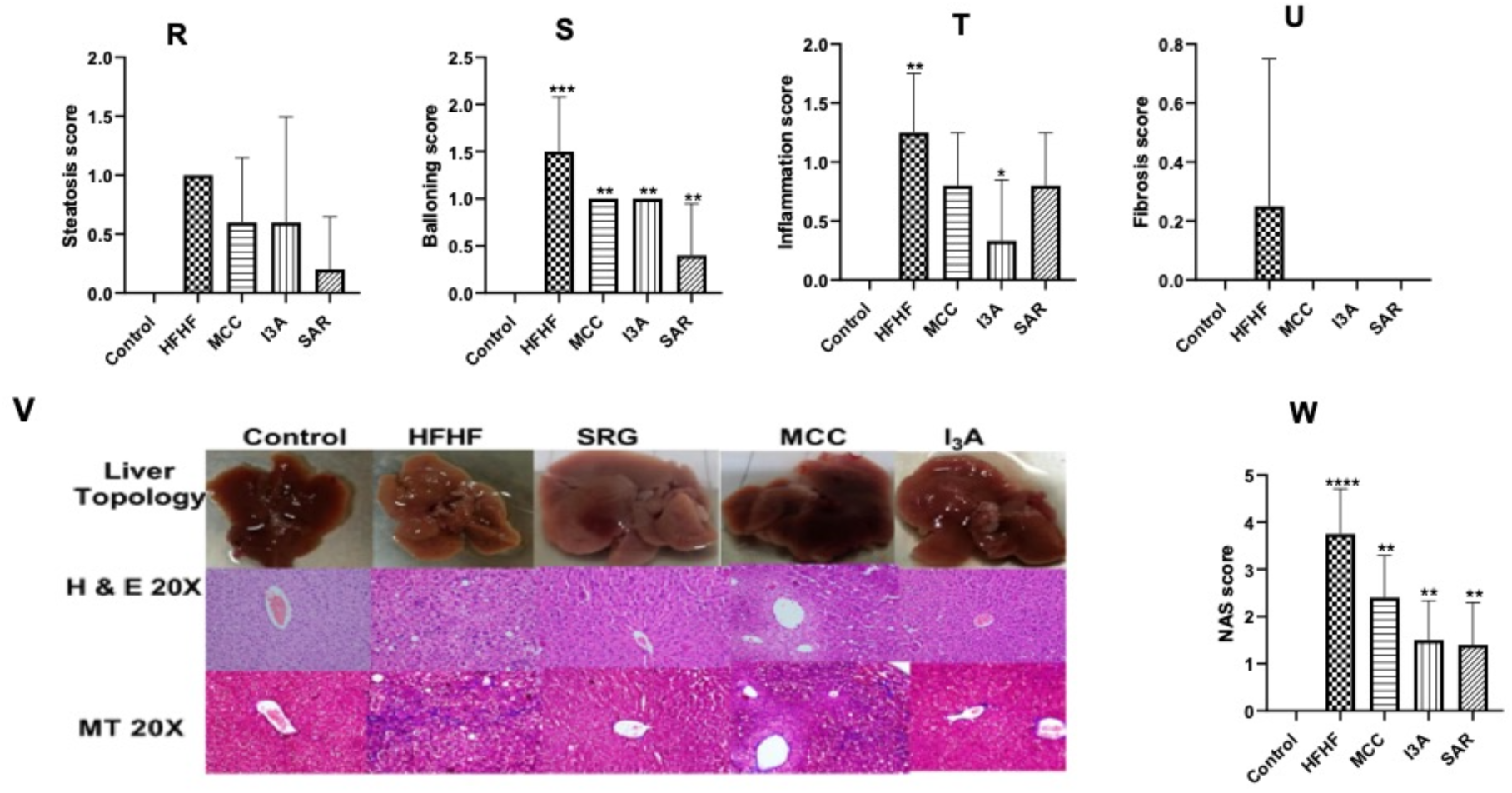
*In vivo* data of I3A-. (A) Change in body weight after 16 weeks of treatment; (B) Body-mass index (BMI); (C) Liver weight. (D) Blood glucose level after fasting (E) AUC curve represents the area under each treatment group (F) Plasma LDL after the end of the treatment period; (G) Plasma Cholesterol after the end of the treatment period;. (H) Plasma TG after the end of the treatment period; (I) Plasma HDL after the end of the treatment period. (J-K) Expression of NLRP3 in HF-HF diet-fed mice, treatment of I3A reduced the expression of NLRP3 and its downstream components IL-1β and IL-18. (J) Hepatic expression of IL-1β (K) Expression of IL-18, data is normalized with mouse *GAPDH* as a housekeeping gene. Quantitative gene expression profiles of inflammatory and fibrotic marker genes from *in vivo* experiments by real time-PCR. (M) TNF-α; (N) IL-6; (P) CXCL1; (Q) TGF-β; (L) α-SMA; (O) COL1A1, data is normalized with mouse *GAPDH* as a housekeeping gene. (R) Steatosis scores from liver histopathology sections (S); Ballooning; (T) Lobular inflammation; (V) NAS scores. (W) Topological images of mice livers from different groups for detecting gross pathological changes (first lane). Representative sections of histopathological analysis of liver tissues (x200 magnification, with H&E stain in the second lane and MT stain in the third lane). Data are represented as Mean ± SEM, n = 6 per group, and statistical analysis consisted of one-way ANOVA followed by Bonferroni’s test (*p < 0.05; ****p < 0.0001 for Control Vs. all other groups.

### 3.6 I3A ameliorates the biochemical parameters in the HF-HF diet-induced model of MASLD

To gain insight into the effect of I3A on HF-HF-induced changes in lipid metabolism, we analysed the biochemical parameters in the blood plasma (Figure 3). The blood glucose level and its AUC (Figures 3D and 3E) were higher in the HF-HF group. A significant reduction in the blood glucose level in mice treated with I3A and Saroglitazar was observed (Figures 3D and 3E). Cholesterol, LDL, and plasma total triglycerides (TG) were significantly increased in mice fed with HF-HF compared with mice subjected to the control group (Figure 3F-3H), and administration with I3A exhibited a reduction in total LDL, cholesterol, and TG as compared with the control HF-HF mice treated with vehicle (Figure 3 F-3H). Our data suggests that treatment with I3A ameliorates key biochemical and metabolic parameters of mice fed with HF-HF diet as compared to vehicle control.

### 3.7 I3A inhibits the mRNA expression levels of IL-1β and IL-18 in mice fed with the HF-HF diet

High expression levels of IL-1β and IL-18 mRNA were observed in the liver tissues of mice in the HF-HF group compared to the control group and administration of I3A significantly diminished the same (Figure 6J and 6K).

### 3.8 I3A attenuates the expression levels of inflammatory and fibrosis markers in the HF-HF diet-induced model of MASLD

The relative gene expression profiling results of mice’s hepatic tissues revealed an increase in the mRNA expression levels of the inflammatory genes, namely TNF-α, IL-6, and CXCL1, in the HF-HF group, which was significantly reduced in mice fed with HF-HF+I3A compared to the disease group (Figure 3M, 3N and 3P). I3A also showed its effect on fibrosis. Treatment with I3A for 18 weeks reduced the mRNA expression of α-SMA, COL1A1, and TGF-β in the liver tissue compared with HF-HF mice treated with vehicle (Figure 3L, 3O and 3Q).

### 3.9 Histopathological Evaluation

The development of steatosis was observed in mice fed with a high-fat, high-fructose (HF-HF) diet for 18 weeks. The livers of mice in the HF-HF group appeared pale, indicating the deposition of lipid, while the livers of mice in the other groups appeared similar to the control group with little discoloration (Figure 3V).

Histopathological examination confirmed the development of the MASLD phenotype in mice in the HF-HF group compared to mice fed with the Chow diet. After 18 weeks of treatment, the livers of mice in the HF-HF group showed significant macrovesicular steatosis, ballooning degeneration, and lobular inflammation (Figure 3V).

The mean MASLD activity score (NAS) in the HF-HF group was 3.75, which was significantly higher compared to the control mice (0), while the HF-HF+SRG and HF-HF+I3A groups had lower mean NAS scores (1.4 & 1.5 respectively) than the disease group (Figure 3W). Early stage fibrosis was observed in one animal in HF-HF group (mean score-0.25) whereas no fibrosis was seen in control, HF-HF+I3A and HF-HF+SRG groups (score-0) (Figure 3U).

Mice in both the HF-HF+I3A and HF-HF+SRG groups, demonstrated a reduction in steatosis, ballooning and inflammation as compared to the HF-HF group. However, the reduction of steatosis and ballooning was much greater with SAR whereas the reduction of inflammation was much greater with I3A. (Figure 3 R,S,T). Our data suggests that the HF-HF diet induces MASLD in mice, and treatment with I3A or SAR reduces the severity of the disease, as evidenced by the improvement in the histopathological scores.

## 4 Discussion

Several studies in the last few years have shown that the gut microbiota plays a crucial role in the regulation of MASLD by synthesizing bacterial metabolites such as short-chain fatty acids, indole derivatives, secondary bile acids, and trimethylamine [20–21]. One study has shown that a leaky gut increases the release of different types of lipids and adipokines, leading to inflammation and lipotoxicity in the liver tissue. In this study, increased serum levels of LPS and gut permeability were found to be associated with the development of MASLD [22]. Administration of probiotics, on the other hand, was found to decrease hepatic steatosis, inflammation, liver injury, and fibrosis in animal models of MASLD [23]. In a recent article by Zhang et al., 2024 [51] fecal metabolomics has been reported as an investigational tool to identify the possible biomarkers for the prognosis and diagnosis of non-obese MASLD. 16S RNA sequencing of fecal DNA from these patients revealed a decrease in the overall richness and variety of the intestinal flora in mice fed a methyl choline deficient diet (MCD). This study reported that g_Tuzzerella, s_Bifidobacterium pseudolongum, and s_Faecalibaculum rodentium were the most common species in non-obese MASLD mice with an overall increase in the Firmicutes: Bacteroidota ratio. Prior to this, Lang S et al. performed 16S rRNA gene sequencing in a cohort of 83 biopsy-proven MASLD patients and 13 patients with non-invasively diagnosed MASLD-cirrhosis to validate the diagnosis of gut microbiota-based approaches over simple non-invasive tools for predicting advanced fibrosis in MASLD. In recent review articles by Chen HT et al., 2020 [52] and Aron-Wisnewsky J et al., 2020 [53], authors have critically evaluated the recent advancements in gut microbiota-targeted therapeutics against MASLD.

Based on these reports, we hypothesized that NLRP3 mediated gut dysbiosis might be playing an important role in the progression of MASLD and modulation of NLRP3 inflammasome activity therefore can be a potential approach to manage MASLD. Further to this we conducted the *in vitro* phenotypic screening of an in-house library of small molecules and natural products and identified, indole-3 acetic acid (I3A), as an inhibitor of NLRP3. I3A was also found to be efficacious in mitigating the key hallmarks of MASLD in mice fed with high fat-high fructose diet in a 22 week study.

Interestingly, I3A has been reported as a gut microbiota-derived metabolite of dietary tryptophan. Tryptophan is an essential amino-acid and the pathways involved in the metabolism of tryptophan have currently been explored as a potential therapeutic target for reducing inflammation in metabolic diseases such as MASLD, obesity and type II diabetes. In another recent publication by Ma et al., 2024 [50], dietary tryptophan was found to significantly increase the relative abundance of Faecalibacterium and Enterococcus. The authors have shown that dietary tryptophan alleviates intestinal inflammatory damage caused by long photoperiod via the inhibition of NLRP3 inflammasome activation. NLRP3 and its family members have been also been previously reported as important contributors to MASLD. Increased levels of NLRP3 mRNA have been reported both in the mouse models of MASLD as well as in patients. Moreover, mice deficient in NLRP3 have been previously shown to be associated with gut dysbiosis [27]. A recent review article by Teunis C et al., 2024 [54] provides a comprehensive analysis of the involvement of three major tryptophan dependent metabolic pathways; such as kynurenine pathway, indole pathway and serotonin/melatonin pathway) which result in metabolites such as kynurenic acid, xanturenic acid, indole-3-propionic acid and serotonin/melatonin. Increased concentrations of the indole pathway metabolites, which are regulated by the gut microbiota, have been found to result in more favourable outcomes for MASLD than the other two pathways. Although Bui TTX; 2019 et al. and Ji Y; 2019, have previously shown the potential of I3A in mitigating MASLD phenotypes in high-fat diet induced models of rat and mouse respectively. However, no data exists to date indicating that the NLRP3 inflammasome pathway is involved in I3A-mediated improvement of MASLD.

In our *in vitro* assay using differentiated THP-1 cells, I3A showed significant inhibition of mRNA levels of IL-1β and IL-18. In addition, I3A also showed inhibition of NLRP3-mediated ASC-SPEC formation in differentiated THP1 cells. In the 3D-spheroid model using a co-culture of Hep-G2 and LX2 (stellate) cells, I3A showed anti-fibrotic activity by inhibiting the levels of palmitic acid-induced COL1A1 and TGF-β. The *in vivo* data reported in this manuscript shows that I3A inhibits the mRNA levels of key signature cytokines of NLRP3 activation, such as IL-1β and IL-18, in the liver tissues of C57BL/6 mice fed with a high-fat-high-fructose (HF-HF) diet. Saroglitazar was used as a positive control in our study. Our data suggests that both Saroglitazar and I3A were effective in reversing the gain in body weight, BMI, and IR in HF-HF-fed mice and also improved the NAS score. I3A-treated mice also appeared healthier than mice fed with the HF-HF diet alone.

The non-availability of an FDA-approved drug for the management of MASLD suggests an urgent need to identify new therapeutic options to manage this disease in a safe and effective manner. Natural products like I3A should therefore be investigated more in detail to obtain molecules with the desired efficacy and fewer side effects compared to existing molecules in the investigational pipeline. Details of the mechanism of action of I3A, as reported in this manuscript, will further help us to conduct dedicated structure-activity-relationship (SAR) studies around the parent molecule and identify molecules with improved efficacy and acceptable safety profile.

## Author contributions

All listed authors participated meaningfully in this study and they have seen and approved the final manuscript.

## Acknowledgements

RT is thankful to Department of Biotechnology, Government of India and BRIC-Translational Health Science and Technology Institute, Faridabad, Haryana, India for providing the financial support to carry out the studies reported in this manuscript.

The authors acknowledge the contributions of Mr. Hari Om for providing technical support in carrying out experiments in mice.

We also acknowledge Small Animal Facility (SAF), THSTI for its services.

## Abbreviations

AIM2: Absent in melanoma 2
ASC: Apoptosis-associated speck-like protein containing a caspase recruitment domain (CARD)
α-SMA: Alpha-smooth muscle actin
CD36: Cluster of differentiation 36 or fatty acid translocase (FAT)
Col1A1: Collagen type 1 α1
FASN: Fatty acid synthase
GasD: Gasdermin D
IL-1β: Interlukin-1 beta
IL-18: Interlukin-18
Casp-1: Caspase-1
IL-6: Interleukin-6
LIPA: Lysosomal acid lipase
MCD: Methionine choline deficient
MASLD: Non-alcoholic fatty liver disease
MASH: Non-alcoholic steatohepatitis
NLRP3: Nucleotide-binding oligomerization domain (NOD), leucine-rich repeat (LRR) (NLRs) containing protein 3
TGF-β: Transforming growth factor beta
TNF-α: Tumor necrosis factor alpha
SCD1: Stearoyl-CoA Desaturase-1

## Supplementary Material

**Table-1.** (Composition of High Fat-High Fructose Diet)

| **Description** | **Ingredients** | **Grams** |
| --- | --- | --- |
| Protein | Casein, Lactic, 30 Mesh | 200.00 g |
| Protein | Cystine, L | 3.00 g |
| Carbohydrate | Fructose | 193.80 g |
| Carbohydrate | Sucrose | 4.00 g |
| Fiber | Solka Floc, FCC200 | 50.00 g |
| Fat | Lard | 245.00 g |
| Fat | Soybean Oil, USP | 25.00 g |
| Mineral | S10026B | 50.00 g |
| Vitamin | Choline Bitartrate | 2.00 g |
| Vitamin | V10001C | 1.00 g |
| Dye | Dye, Yellow FD&C #5, Alum. Lake 35-42% | 0.03 g |
| Dye | Dye, Blue FD&C #1, Alum. Lake 35-42% | 0.03 g |

**Table-1A.** (Mineral S10026B Composition)

| **Description** | **Ingredients** | **Grams** |
| --- | --- | --- |
| Carbohydrate | Sucrose, Fine Granulated | 179.82 g |
| Mineral | Potassium Citrate, Monohydrate | 330.00 g |
| Mineral | Calcium Phosphate, Dibasic | 260.00 g |
| Mineral | Calcium Carbonate, Light, USP | 110.00 g |
| Mineral | Sodium Chloride | 51.80 g |
| Mineral | Magnesium Sulfate, Heptahydrate | 51.52 g |
| Mineral | Magnesium Oxide, Heavy, DC USP | 8.38 g |
| Mineral | Ferric Citrate | 4.20 g |
| Mineral | Manganese Carbonate Hydrate | 2.45 g |
| Mineral | Zinc Carbonate | 1.12 g |
| Mineral | Chromium Potassium Sulfate | 0.39 g |
| Mineral | Copper Carbonate | 0.21 g |
| Mineral | Ammonium Molybdate Tetrahydrate | 0.06 g |
| Mineral | Sodium Fluoride | 0.04 g |
| Mineral | Sodium Selenite | 0.01 g |
| Mineral | Potassium Iodate | 0.01 g |
|  | Total: | 1000.00 g |

**Table-1B.**
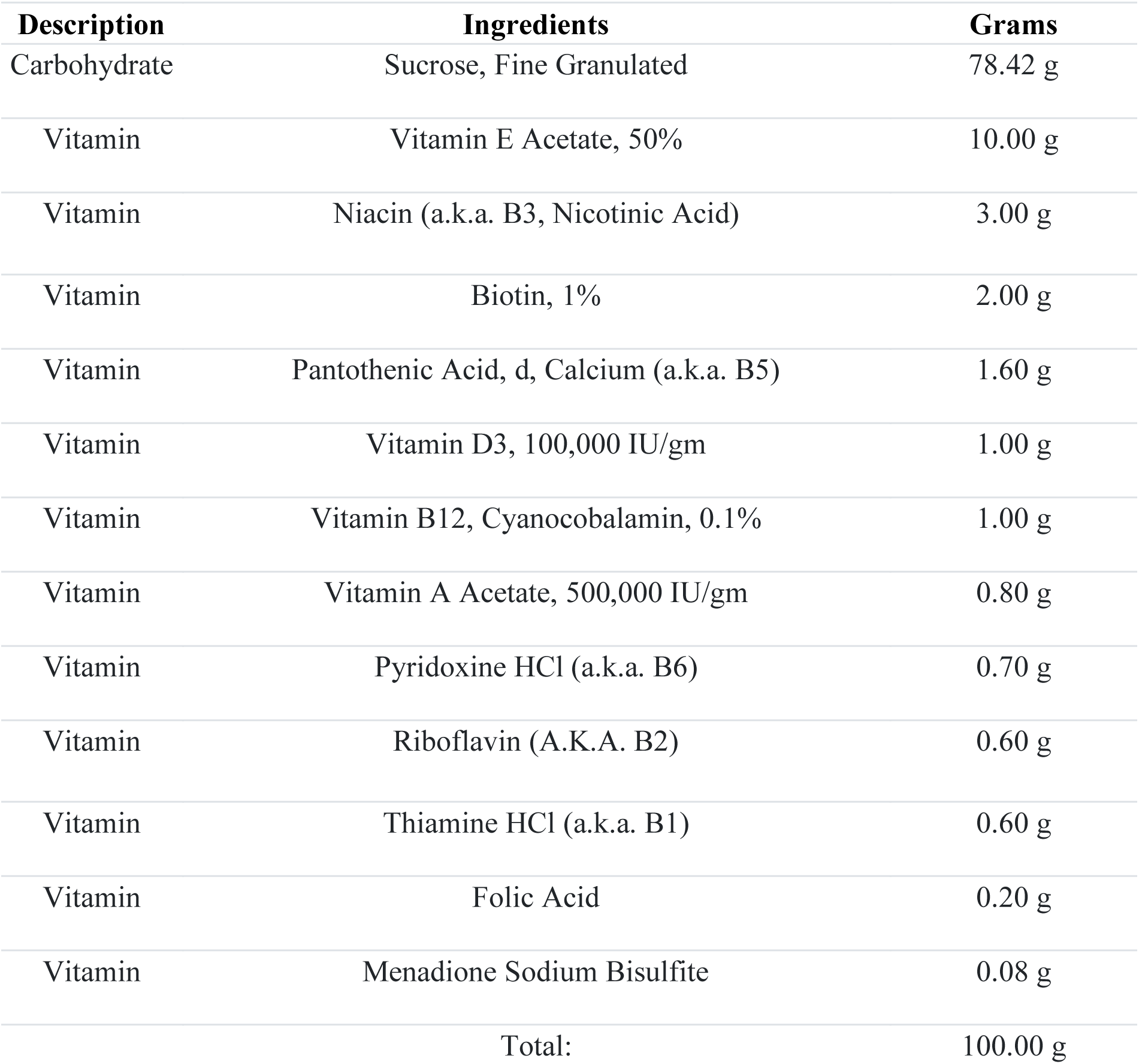
Vitamin V10001C Composition

**Table-2.**
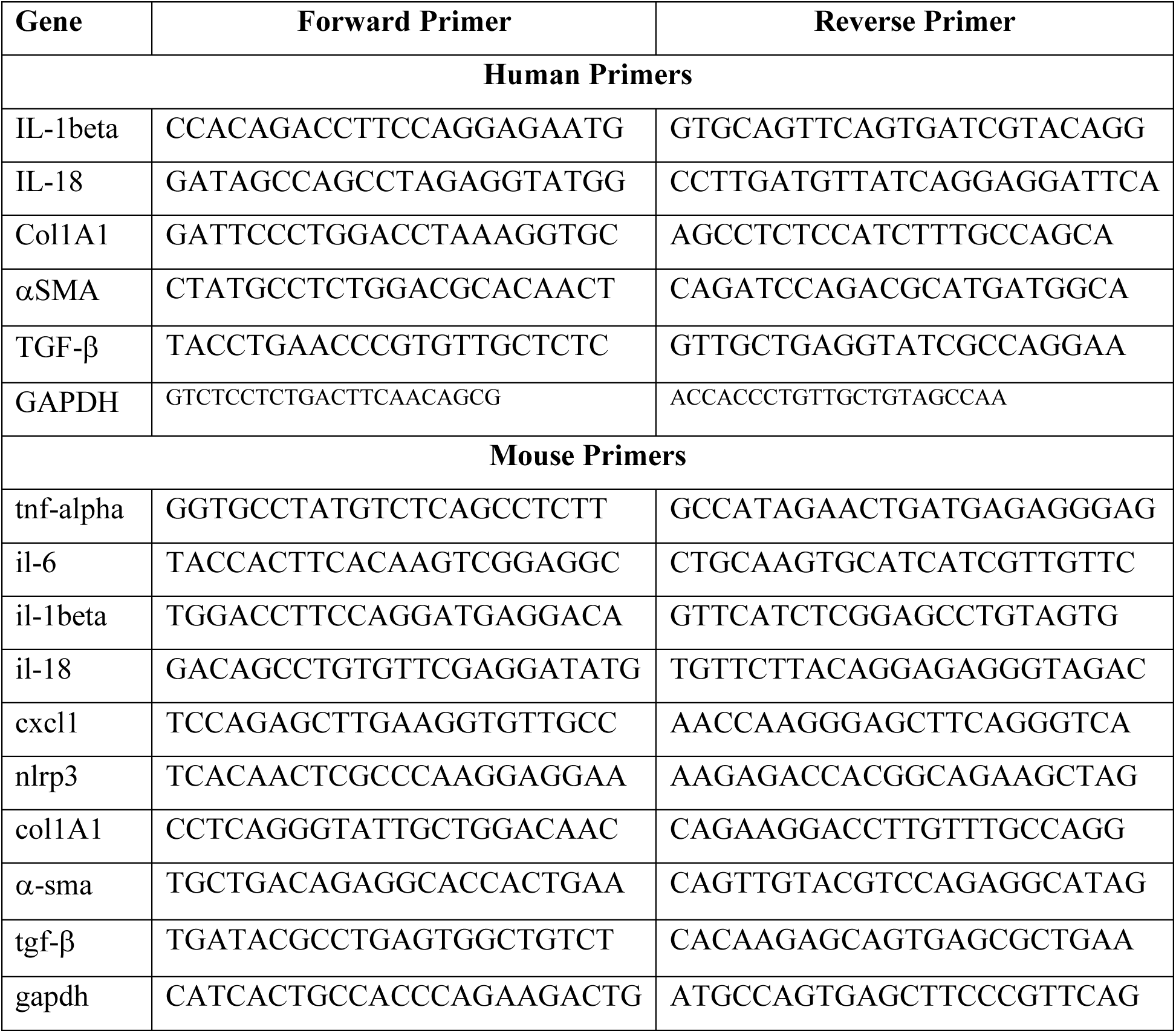
List of primers

## References

1. Norheim, F.; Hui, S.T.; Kulahcioglu, E.; Mehrabian, M.; Cantor, R.M.; Pan, C.; Parks, B.W.; Lusis, A.J. Genetic and hormonal control of hepatic steatosis in female and male mice. J. Lipid Res. 2017, 58, 178–187.

2. Abd El-Kader, S.M.; El-Den Ashmawy, E.M. Non-alcoholic fatty liver disease: The diagnosis and management. World J. Hepatol. 2015, 7, 846–858.

3. Younossi, Z.; Anstee, Q.M.; Marietti, M.; Hardy, T.; Henry, L.; Eslam, M.; George, J.; Bugianesi, E. Global burden of MASLD and MASH: Trends, predictions, risk factors and prevention. Nat. Rev. Gastroenterol. Hepatol. 2018, 15, 11–20.

4. Rau, M.; Schilling, A.K.; Meertens, J.; Hering, I.; Weiss, J.; Jurowich, C.; Kudlich, T.; Hermanns, H.M.; Bantel, H.; Beyersdorf, N.;, et al. Progression from Nonalcoholic Fatty Liver to Nonalcoholic Steatohepatitis Is Marked by a Higher Frequency of Th17 Cells in the Liver and an Increased Th17/Resting Regulatory T Cell Ratio in Peripheral Blood and in the Liver. J. Immunol. 2016, 196, 97–105.

5. Takahashi, Y.; Fukusato, T. Histopathology of nonalcoholic fatty liver disease/nonalcoholic steatohepatitis. World J. Gastroenterol. 2014, 20, 15539–15548.

6. Buzzetti, E.; Pinzani, M.; Tsochatzis, E.A. The multiple-hit pathogenesis of metabolic-dysfunction associated steatotic liver disease(MASLD). Metabolism 2016, 65, 1038–1048.

7. Fang, Y.L.; Chen, H.; Wang, C.L.; Liang, L. Pathogenesis of metabolic-dysfunction associated steatotic liver diseasein children and adolescence: From “two hit theory” to “multiple hit model”. World J. Gastroenterol. 2018, 24, 2974–2983.

8. Ipsen, D.H.; Lykkesfeldt, J.; Tveden-Nyborg, P. Molecular mechanisms of hepatic lipid accumulation in non-alcoholic fatty liver disease. Cell Mol. Life Sci. 2018, 75, 3313–3327.

9. Li, Z.Z.; Berk, M.; McIntyre, T.M.; Feldstein, A.E. Hepatic lipid partitioning and liver damage in nonalcoholic fatty liver disease: Role of stearoyl-CoA desaturase. J. Biol. Chem. 2009, 284, 5637–5644.

10. Masarone, M.; Rosato, V.; Dallio, M.; Gravina, A.G.; Aglitti, A.; Loguercio, C.; Federico, A.; Persico, M. Role of Oxidative Stress in Pathophysiology of Nonalcoholic Fatty Liver Disease. Oxid. Med. Cell. Longev. 2018, 2018, 9547613.

11. Li, S.; Hong, M.; Tan, H.Y.; Wang, N.; Feng, Y. Insights into the Role and Interdependence of Oxidative Stress and Inflammation in Liver Diseases. Oxid. Med. Cell. Longev. 2016, 2016, 4234061.

12. Knorr, J.; Wree, A.; Tacke, F., Feldstein, A.E. The NLRP3 Inflammasome in Alcoholic and Nonalcoholic Steatohepatitis. Semin Liver Dis. 2020 Aug;40(3):298–306.

13. Yu, L.; Hong, W.; Lu, S.; Li, Y.; Guan, Y.; Weng, X.; Feng, Z. The NLRP3 Inflammasome in NonAlcoholic Fatty Liver Disease and Steatohepatitis: The rapeutic Targets and Treatment. Front Pharmacol. 2022;13:780496.

14. Benetti, E.; Mastrocola, R.; Vitarelli, G.; Cutrin, J. C.; Nigro, D.; Chiazza, F.; Mayoux, E.; Collino, M.; Fantozzi, R. Empagliflozin Protects against Diet-Induced NLRP-3 Inflammasome Activation and Lipid Accumulation., J Pharmacol Exp Ther. 2016, 359, 45–53.

15. Mridha, A. R.; Wree, A.; Robertson, A. A. B.; Yeh, M. M.; Johnson, C. D.; Van Rooyen, D. M.; Haczeyni, F.; Teoh, N. C.; Savard, C.; Ioannou, G. N.; Masters, S. L.; Schroder, K.; Cooper, M. A.; Feldstein, A. E.; Farrell, G. C. NLRP3 inflammasome blockade reduces liver inflammation and fibrosis in experimental MASH in mice., J Hepatol. 2017, 66, 1037–1046.

16. Wree, A.; McGeough, M.D.; Peña, C.A.; Schlattjan, M.; Li, H.; Inzaugarat, M.E.; Messer, K.; Canbay, A.; Hoffman, H.M.; Feldstein, A.E. NLRP3 inflammasome activation is required for fibrosis development in MASLD. J. Mol. Med. 2014, 92, 1069–1082.

17. Zhu, X.; Lin, X.; Zhang, P.; Liu, Y.; Ling, W.; Guo, H. Upregulated NLRP3 inflammasome activation is attenuated by anthocyanins in patients with nonalcoholic fatty liver disease: A case-control and an intervention study Clin Res Hepatol Gastroenterol, 2022;46(4):101843. doi: 10.1016/j.clinre.2021.101843

18. Hubbard, T.D.; Murray, I.A.; Perdew, G.H. Indole and Tryptophan Metabolism: Endogenous and Dietary Routes to Ah Receptor Activation. Drug Metab. Dispos. 2015, 43, 1522–1535.

19. Krishnan, S.; Ding, Y.; Saedi, N.; Choi, M.; Sridharan, G.V.; Sherr, D.H.; Yarmush, M.L.; Alaniz, R.C.; Jayaraman, A.; Lee, K. Gut Microbiota-Derived Tryptophan Metabolites Modulate Inflammatory Response in Hepatocytes and Macrophages. Cell Rep. 2018, 23, 1099–1111.

20. Zhu, Y.; Liu, H.; Zhang, M.; Guo, G.L. Fatty liver diseases, bile acids, and FXR. Acta Pharm. Sin. B 2016, 6, 409–412.

21. Juarez-Hernandez, E.; Chavez-Tapia, N.C.; Uribe, M.; Barbero-Becerra, V.J. Role of bioactive fatty acids in nonalcoholic fatty liver disease. Nutr. J. 2016, 15, 72.

22. Farhadi, A.; Gundlapalli, S.; Shaikh, M.;, et al. Susceptibility to gut leakiness: a possible mechanism for endotoxaemia in non-alcoholic steatohepatitis. Liver International. 2008;28(7):1026–1033. doi: 10.1111/j.1478-3231.2008.01723.x.

23. Velayudham, A.; Dolganiuc, A.; Ellis, M.;, et al. VSL#3 probiotic treatment attenuates fibrosis without changes in steatohepatitis in a diet-induced nonalcoholic steatohepatitis model in mice. Hepatology. 2009;49(3):989–997. doi: 10.1002/hep.22711.

24. Roager, H.M.; Licht, T.R. Microbial tryptophan catabolites in health and disease. Nat. Commun. 2018, 9, 3294.

25. Rath, S.; Heidrich, B.; Pieper, D.H.; Vital, M. Uncovering the trimethylamine-producing bacteria of the human gut microbiota. Microbiome. 2017, 5, 54.

26. Xie, N.; Cui, Y.; Yin, Y.-N.;, et al. Effects of two Lactobacillus strains on lipid metabolism and intestinal microflora in rats fed a high-cholesterol diet. BMC Complementary and Alternative Medicine. 2011;11, article 53 doi: 10.1186/1472-6882-11-53.

27. Pan, H.; Jian, Y.; Wang, F.; Yu, S.; Guo, J.; Kan, J.; Guo, W. NLRP3 and Gut Microbiota Homeostasis: Progress in Research. Cells. 2022, 11(23), 3758. doi: 10.3390/cells11233758.

28. Lamkanfi, M. Emerging inflammasome effector mechanisms. Nat Rev Immunol. 2011, 11, 213–220.

29. Arnao, M.B.; Sanchez-Bravo, J.; Acosta, M. Indole-3-carbinol as a scavenger of free radicals. Biochem. Mol. Biol. Int. 1996, 39, 1125–1134.

30. Kim, D.; Kim, H.; Kim, K.; Roh, S. The Protective Effect of Indole-3-Acetic Acid (I3A) on H2O2 -Damaged Human Dental Pulp Stem Cells Is Mediated by the AKT Pathway and Involves Increased Expression of the Transcription Factor Nuclear Factor-Erythroid 2-Related Factor 2 (Nrf2 and Its Downstream Target Heme Oxygenase 1 (HO-1). Oxid. Med. Cell. Longev. 2017, 2017, 8639485.

31. Ji, Y.; Gao, Y.; Chen, H.; Yin, Y.; Zhang, W. Indole-3-Acetic Acid Alleviates Non alcoholic Fatty Liver Disease in Mice via Attenuation of Hepatic Lipogenesis, and Oxidative and Inflammatory Stress., Nutrients 2019, 11, 2062.

32. Bui, T.T.X.; Nguyen, D.T.K.H.; Cao, D.D.T.; Hoang, N.T.M.; Ly, V.V.N.; Nguyen, H.N.T.Q.; et al. Therapeutic effect of indole-3 acetic acid on non-alcoholic fatty liver disease in rats fed a high fat diet [published correction appears in J Physiol Biochem. 2019 Jul;75(3):555]. J Physiol Biochem 2018;74(4). doi:10.1007/s13105-018-0654-5.

33. Pingitore, P.;, et al., Human Multilineage 3D Spheroids as a Model of Liver Steatosis and 844 Fibrosis. Int J Mol Sci, 2019. 20(7).

34. Jain, M.R.;, et al., Dual PPARalpha/gamma agonist saroglitazar improves liver 860 histopathology and biochemistry in experimental MASH models. Liver Int, 2018. 38(6): p. 861 1084–1094.

35. Kumari D, Gautam J, Sharma V, Gupta S, Sarkar S, Jana P, Singhal V, Babele P, Kamboj P, Bajpai S, Tandon R, Kumar Y, Dikshit M. Effect of herbal extracts and Saroglitazar on high-fat diet-induced obesity, insulin resistance, dyslipidemia, and hepatic lipidome in C57BL/6J mice, Heliyon (2023).

36. Bank RPD. RCSB PDB: Homepage. [cited 7 Oct 2022]. Available: https://www.rcsb.org/

37. Kumari, A.; Mittal, L.; Srivastava, M.; Pathak, D.P.; Asthana, S. Deciphering the Structural Determinants Critical in Attaining the FXR Partial Agonism. J Phys Chem B. 2023;127: 465–485.

38. Mittal, L.; Srivastava, M.; Kumari, A.; Tonk, R.K.; Awasthi, A.; Asthana, S. Interplay among Structural Stability, Plasticity, and Energetics Determined by Conformational Attuning of Flexible Loops in PD-1. J Chem Inf Model. 2021;61: 358–384.

39. Dekker, C.; Mattes, H.; Wright, M.; Boettcher, A.; Hinniger, A.; Hughes, N.;, et al. Crystal Structure of NLRP3 NACHT Domain With an Inhibitor Defines Mechanism of Inflammasome Inhibition. J Mol Biol. 2021;433: 167309.

40. Srivastava, M.; Mittal, L.; Kumari, A.; Agrahari, A.K.; Singh, M.; Mathur, R.;, et al. Characterizing (un)binding mechanism of USP7 inhibitors to unravel the cause of enhanced binding potencies at allosteric checkpoint. Protein Sci. 2022;31: e4398.

41. Sastry, G.M.; Adzhigirey, M.; Day, T.; Annabhimoju, R.; Sherman, W. Protein and ligand preparation: parameters, protocols, and influence on virtual screening enrichments. J Comput Aided Mol Des. 2013;27: 221–234.

42. Panwar, S.; Kumari, A.; Kumar, H.; Tiwari, A.K.; Tripathi, P.; Asthana, S. Structure-based virtual screening, molecular dynamics simulation and evaluation to identify inhibitors against NAMPT. J Biomol Struct Dyn. 2022;40: 10332–10344.

43. Mittal, L.; Kumari, A.; Suri, C.; Bhattacharya, S.; Asthana, S. Insights into structural dynamics of allosteric binding sites in HCV RNA-dependent RNA polymerase. J Biomol Struct Dyn. 2020;38: 1612–1625.

44. Kumari, A.; Mittal, L.; Srivastava, M.; Asthana, S. Binding mode characterization of 13b in the monomeric and dimeric states of SARS-CoV-2 main protease using molecular dynamics simulations. J Biomol Struct Dyn. 2022;40: 9287–9305.

45. Mittal, L.; Tonk, R.K.; Awasthi, A.; Asthana, S. Targeting cryptic-orthosteric site of PD-L1 for inhibitor identification using structure-guided approach. Arch Biochem Biophys. 2021;713: 109059.

46. Friesner, R.A.; Banks, J.L.; Murphy, R.B.; Halgren, T.A.; Klicic, J.J.; Mainz, D.T.;, et al. Glide: a new approach for rapid, accurate docking and scoring. 1. Method and assessment of docking accuracy. J Med Chem. 2004;47: 1739–1749.

47. Genheden, S.; Ryde, U. The MM/PBSA and MM/GBSA methods to estimate ligand-binding affinities. Expert Opin Drug Discov. 2015;10: 449–461.

48. Singh, M.; Srivastava, M.; Wakode, S.R..; Asthana S. Elucidation of Structural Determinants Delineates the Residues Playing Key Roles in Differential Dynamics and Selective Inhibition of Sirt1-3. J Chem Inf Model. 2021;61: 1105–1124.

49. Interplay among Structural Stability, Plasticity and Energetics Determined by Conformational Attuning of Flexible Loops in PD1. doi:10.1021/acs.jcim.0c01080.s001

50. Ma D, Zhang S, Zhang M, Feng J. Dietary tryptophan alleviates intestinal inflammation caused by long photoperiod via gut microbiota derived tryptophan metabolites-NLRP3 pathway in broiler chickens. Poult Sci. 2024 Apr;103(4):103509. doi: 10.1016/j.psj.2024.103509. Epub 2024 Jan 29. PMID: 38387289; PMCID: PMC10900804.

51. Zhang W, Cheng W, Li J, Huang Z, Lin H, Zhang W. New aspects characterizing non-obese MASLD by the analysis of the intestinal flora and metabolites using a mouse model. mSystems. 2024 Mar 19;9(3):e0102723. doi: 10.1128/msystems.01027-23. Epub 2024 Feb 29. PMID: 38421203; PMCID: PMC1094948.

52. Chen HT, Huang HL, Li YQ, Xu HM, Zhou YJ. Therapeutic advances in non-alcoholic fatty liver disease: A microbiota-centered view. World J Gastroenterol. 2020 Apr 28;26(16):1901–1911. doi: 10.3748/wjg.v26.i16.1901. PMID: 32390701; PMCID: PMC7201149.

53. Aron-Wisnewsky J, Warmbrunn MV, Nieuwdorp M, Clément K. Nonalcoholic Fatty Liver Disease: Modulating Gut Microbiota to Improve Severity? Gastroenterology. 2020 May;158(7):1881–1898. doi: 10.1053/j.gastro.2020.01.049. Epub 2020 Feb 8. PMID: 32044317.

54. Teunis C, Nieuwdorp M, Hanssen N. Interactions between Tryptophan Metabolism, the Gut Microbiome and the Immune System as Potential Drivers of Metabolic-dysfunction associated steatotic liver disease(MASLD) and Metabolic Diseases. Metabolites. 2022 Jun 2;12(6):514. doi: 10.3390/metabo12060514. PMID: 35736447; PMCID: PMC9227929.

